# Contralateral delay activity during dynamic spatial updates in working memory

**DOI:** 10.1101/2024.02.27.582343

**Authors:** Charles Chernik, Ronald van den Berg, Mikael Lundqvist

## Abstract

Working Memory (WM) enables us to maintain and directly manipulate mental representations, yet we know little about the neural implementation of this privileged online format. We recorded electroencephalography data as human subjects engaged in a task requiring continuous updates to the locations of objects retained in WM as well as in a visually identical task with WM demands removed. Analysis of neural data suggested WM-related contralateral delay activity reversed polarity as objects held in WM moved to the opposite hemifield. This was partially but not fully explained by visual attention demands. Thus, the cortical location of activity related to both attention and WM was updated to meet behavioral demands as the spatial location of remembered objects changed.

## Introduction

A fundamental aspect of cognition is our ability to hold and mentally manipulate information in an online state in working memory (WM)^1,2^. Both the format of retention^3–5^ and the actual manipulation of WM information^6–10^ are highly studied topics. Despite this, we know relatively little about the transformations of memory representations that underlie such manipulations^11^. This knowledge gap may in part be due to WM representations typically being studied under relatively static settings, while in real life WM representations are continuously updated to support behavior^12,13^.

Spiking and local-field potential data from monkeys suggest that the representation of visual objects transfers from one prefrontal hemisphere to the other following saccades that move the location of an earlier presented stimulus to the opposite hemifield^12^. Likewise, gaze bias in humans, a behavioral correlate of the location of retained WM objects, shifts as subjects turn their heads^13^. These findings suggest that WM object representations transfer across cortex when humans mentally update their remembered locations, but electrophysiological evidence for this is lacking. Here, we aim to bridge that gap by using a novel experimental paradigm combined with electroencephalogram (EEG) measures.

We tracked the cortical locations of WM representations using Contralateral Delay Activity (CDA), an electrophysiological measure of WM representations measured via EEG^14,15^. This measure reflects the difference in brain activity between the electrodes contralateral and ipsilateral to the side where the remembered items were presented. It correlates with the number of relevant but not irrelevant visual WM objects, and is often used as a correlate of lateral WM load^14^. Prior work has shown that CDA is sensitive to the selection of a subset of particular items after initial encoding^16^, as well as subsequent encoding of additional items^17^. Due to its lateralized nature, CDA offers a way to test whether memory representations are spatially transferred within the cortex when remembered stimuli are mentally relocated from one visual hemifield to the other. Here, we have tested this by recording EEG while participants performed a WM task that required them to mentally update the locations of two remembered items across visual hemifields. Our main hypothesis was that when the remembered locations of stimuli move across visual hemifields, the underlying neural representations likewise relocate across hemispheres, reversing the sign of the CDA signal.

## Results

### Experiment 1: Working memory

We recorded EEG from healthy humans during a delayed-estimation task^18^ that involved mental updating of locations of memorized stimuli after they disappeared from the screen (Figure 1a). Each trial began with a central fixation, followed by a cue indicating on which side (left or right of the fixation point) the to-be-remembered objects would appear. Four differently colored discs (two on each side) were then shown, with the two non-cued items serving to balance visual inputs in the two hemispheres and ensuring activity differences were due to memory-related processes. After 500 ms, the four discs were replaced by colorless outlines that started moving 1 s later along an (invisible) circle (180°/s, clockwise or counterclockwise, 0-360 degrees, with 13 possible distances; Figure 1b). Finally, 3 s after movement onset, one of the outlines was highlighted and the subject was prompted to provide their estimate of its remembered color through a mouse click on a color wheel. Importantly, since the direction and degree of movement were unpredictable, accurate performance was possible only if the subjects mentally updated item positions in WM in real time.

**Figure 1.**
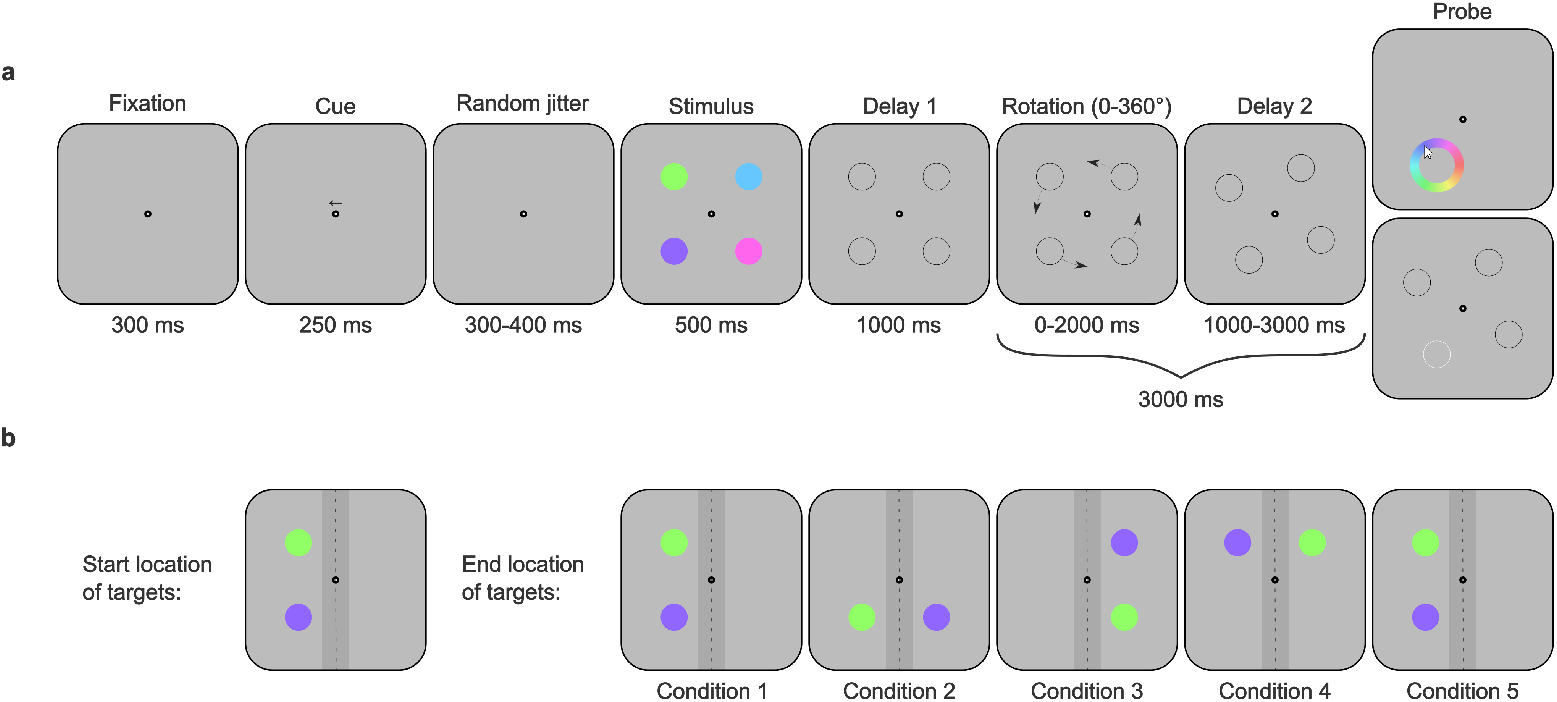
Task design of the experiments. **a.** Trial structure. Each trial started with a fixation period of 300 ms, followed by a cue indicating the to-be-attended side (left or right). After a random jitter of 300-400 ms, a memory array of four colored discs appeared, with two on each side of the fixation point. After 500 ms, the colored discs were replaced with black outlines to indicate their locations. After 1000 ms the outlines simultaneously started moving along an invisible circle, changing their positions relative to the midline by 0, 30, …, or 360 degrees (13 possible alternatives, with randomized sign, 180 deg/s). Once the movement was complete, the outlines remained in their new positions for at least 1 s. Then, in Experiment 1 (top), all the outlines disappeared, and a color wheel was presented at the post-movement location of one of the memorized discs, prompting recall. The response was submitted via a mouse click anywhere on the color wheel. In Experiment 2 (bottom), the outlines stayed in place, with one of them changing its color to white. The subject indicated through a keyboard button press whether the white outline corresponded to one of the tracked discs. **b**. Schematic illustration of the five experimental conditions, defined by the laterality of the post-movement stimulus locations: Condition 1 – both discs remained in the original hemifield; Condition 2 – one of the discs transferred to the other hemifield; Condition 3 – both discs transferred to the other hemifield; Condition 4 – both discs transferred to the other hemifield and then one of them returned to the original hemifield; and Condition 5 – both discs transferred to the other hemifield and then returned to the original one. While there were 13 possible rotation angles, each of them fell in one of these five conditions. The dark gray region represents the area along the midline (width = 1.6°) where the discs could not end up after the movement, ensuring lateralization.

The response error with respect to the target item gradually increased with larger movement angles (Figure 2a, green), while the error with respect to the non-target item (the relevant item not probed) decreased (Figure 2a, purple). This suggests that subjects tended to confuse the two retained colors (swapping errors^19^) more often with larger movements. This was corroborated by fitting the data with an equal-precision model with non-target responses^20^, which explained the performance degradation through increasing swapping rates at larger movements, while memory precision of the two items remained stable (Figure 2b). Importantly, even at the largest movement angle subjects performed accurately: estimated swap rates were relatively low (around 10%) and memory precision was high (parameter log(*κ*) was on average around 2, which corresponds to an average memory error of 16.8° on non-swap trials). Hence, they must have been updating item position in WM in real time.

**Figure 2.**
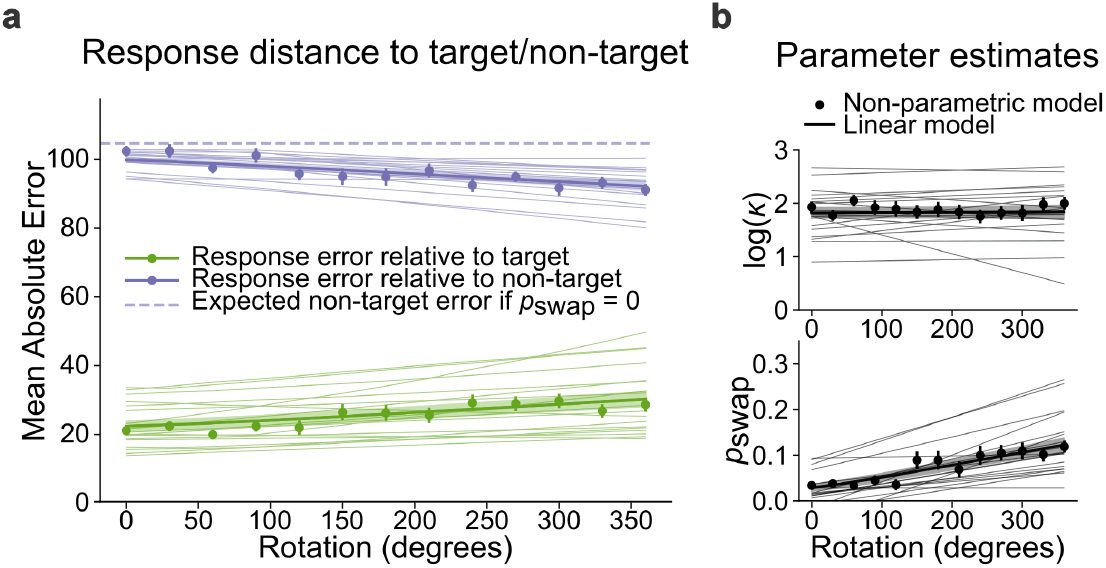
Effect of stimulus movement on response accuracy. **a.** Subject-averaged response errors as a function of movement angle. The error relative to the target (green markers) gradually increased with movement angle (Pearson’s *r*=.88, *p*<.001), while the error relative to the non-target item (purple markers) decreased (*r*=−.89, *p*<.001). These trends were well accounted for by a model in which memory precision and swap rate are linearly related to motion angle (thin lines: estimates for individual subjects; thick lines: averages). Shaded areas and error bars represent one SEM. **b**. Estimated effect of movement angle on memory precision (top) and swap rates (bottom) in the linear model (lines) and in a non-parametric model (markers). While swap rate increased (*r*=.92, *p*<.001), there was no evidence that memory precision was affected by motion angle (*r*=−.02, *p*=.93).

Next, we computed the CDA from the EEG data to investigate the neural underpinning of this mental updating. Under the hypothesis that mental updating of item locations caused WM representations to physically transfer across the cortex (Model 1) both the magnitude and sign of the CDA were predicted to change in specific ways as the empty discs moved across visual hemifields (Figure 3a): right before motion onset, the CDA should be negative, as in previous studies using stationary items; in trials where both discs stayed in the original hemifield (Condition 1), the CDA should remain negative; if one disc moved over to the other hemifield (Condition 2), the CDA should disappear, because the two WM objects would be in opposite hemispheres, eliminating the lateralized imbalance that drives the CDA; if both discs crossed the midline (Condition 3), the CDA should change sign to positive; if one disc then moved back into the original hemifield, the CDA should again disappear (Condition 4); finally, in the trials where both discs moved full circle (Condition 5), the post-motion CDA should be negative again, as if no rotation had taken place. An alternative hypothesis is that the neural representations of WM objects remain anchored to their original encoding locations, regardless of subsequent mental updates of spatial positions (Model 2). This model predicts that the sign of the CDA remains negative throughout any post-encoding updates of locations, with a magnitude that is either stable or may gradually decrease to reflect decaying memory quality. A final hypothesis is that mental movement turns the neural representations into a new format that is not tracked by CDA (Model 3). This model predicts that cross-hemispheric movements cause CDA (possibly apart from Condition 1) to disappear and not come back.

**Figure 3.**
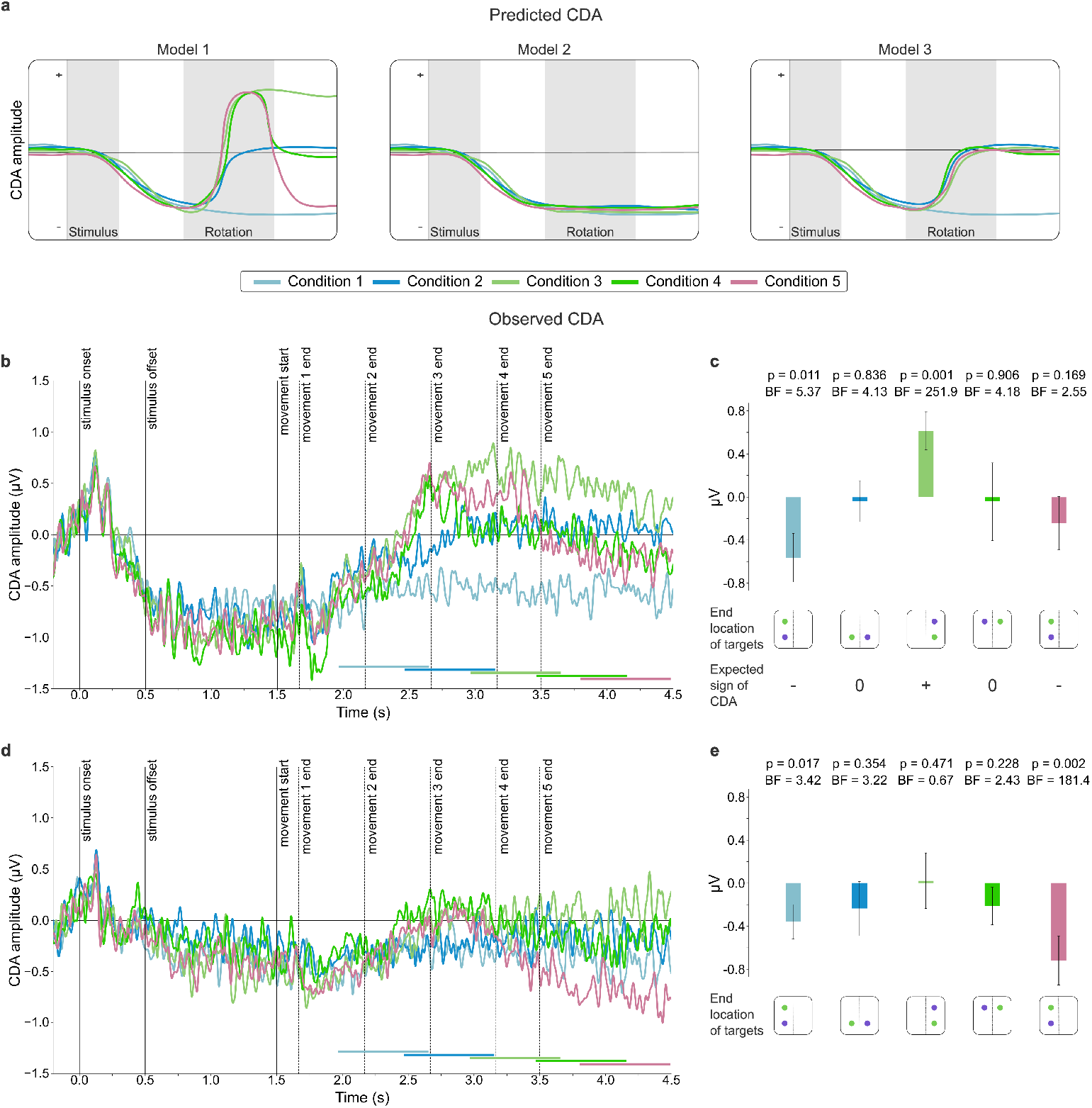
CDA following movement of WM objects/attentional tracking. **a.** Cartoon of model predictions of how the CDA changes following the mental relocation of the discs. In the main model, Model 1, the CDA changes depending on the hemifield the discs end up in post rotation (condition 1-5). For condition 1 and 5 the discs end up in the initial hemifield and the CDA should be negative. For condition 2 and 4, there is one disc in each hemifield and there should be no CDA. For condition 3, both discs end up in the opposite hemifield and the CDA should be positive. As alternatives, Model 2 predicts that the neural representations of the disks remain in their original locations (meaning CDA remains unchanged) and Model 3 predicts the CDA disappears completely after the update of WM contents occurs (meaning the CDA only remains when both discs stay in the initial hemifield). **b**. Measured CDA per condition in Experiment 1. Dotted vertical lines denote the latest possible time when movement ended per condition. Colored horizontal bars denote 300-1000 ms post movement per condition, used to measure the CDA statistically. **c**. *Top:* average CDA post movement per condition in Experiment 1, taken from the time periods indicated in **b**. Error bars denote SEM. The p-values were obtained from t-tests on the null hypothesis of CDA being 0. In the conditions where Model 1 predicted a specific effect direction, one-sided tests were used. The BFs were obtained from Bayesian t-tests and quantify the evidence in favor of Model 1 (see Methods). *Middle:* Cartoon demonstrating the end location of the targets per condition (assuming both discs started on the left). *Bottom:* sign of the CDA predicted by Model 1. **d**. Measured CDA per condition in Experiment 2. **e**. Average CDA post movement per condition in Experiment 2.

Qualitatively, our data closely aligned with the predictions of Model 1, with the CDA starting off negative, reversing sign as WM objects transitioned into the opposite hemifield and then again when they moved back into the original hemifield (Figure 3b). To test the predictions of Model 1 quantitatively, we performed t-tests on the time-averaged CDA in each of the five conditions, in a 700 ms window shortly after movement ended in each condition (Figure 3b, horizontal bars). For all 5 conditions, the Bayes Factors (BF) supported the prediction of Model 1 (Figure 3c, n=19 in all tests), though for Condition 5 the evidence was anecdotal (BF = 2.55). Similarly, the p-values were in line with the model’s predictions for all conditions except Condition 5. These results were robust against changes in how the CDA time windows were defined (Supplemental Figure 1 and text). In the three conditions where Models 1 and 2 differed in their predictions, the data favored Model 1, with BFs of 3.57 (Condition 2), 471.7 (Condition 3), and 3.84 (Condition 4). In the comparison between Models 1 and 3, Model 1 was clearly favored in Condition 3 (BF = 32.1), but the evidence was weakly in favor of Model 3 in Condition 5 (BF = 0.60). Finally, we fitted Bayesian mixed-effects models to compare the three Models using all five conditions at once. This analysis again favored Model 1, capturing the predicted CDA across conditions with an estimated model weight of 59%, versus 25% for Model 3 and 16% for Model 2.

### Experiment 2: Attentional tracking

While the CDA is widely regarded as a neural marker of working memory (WM) storage ^15,21,22^, some studies suggest that it is also sensitive to attentional processes ^23–25^. To test to what extent the patterns observed in Experiment 1 reflect attentional processes rather than WM retention, we conducted another experiment with the same sequence of stimuli as in Experiment 1, but with a task that did not require storing them in WM. In Experiment 2, participants were instructed to ignore the colors and simply keep track of the positions of the two cued discs. At probe, instead of recalling the color of the highlighted outline, they indicated whether it was one of the two tracked discs. If the cortical transfer suggested by the CDA patterns in Experiment 1 was primarily a marker of attentional tracking, we expected the pattern in Experiment 2 to be similar, both qualitatively and quantitatively. However, if cortical transfer of WM contents was an important driver of the patterns observed in Experiment 1, then the CDA and sign reversals should be less pronounced in Experiment 2.

As this was a simple task, the behavioral performance was unsurprisingly good (accuracy = 94.2±5.1%, M ± SD). Since the task in Experiment 2 did not involve WM encoding, we first checked if the stimuli evoked a CDA signal at all, by performing a Bayesian t-test on the CDA in the 700 ms window before the movement (this part of the trial was identical across the five conditions). Evidence for a negative CDA was observed (BF = 22.7), albeit of much weaker strength than in Experiment 1 (BF = 141.03). Hence, the CDA in our paradigm appears to reflect a mixture of attentional and WM processes, with attention alone producing a weaker and less consistent signal.

Next, we computed and analyzed the post-movement CDA the same way as before. Qualitatively, the observed pattern was comparable to the one in Experiment 1, yet there were marked quantitative differences (Figure 3d). Notably, unlike in Experiment 1, no polarity reversal from negative to positive was observed in Condition 3 (Figure 3e). Bayesian mixed-effect modelling gave 47%, 45%, and 8% weight to Models 1, 2, and 3, respectively.

In summary, Experiment 2 showed that although attentional tracking alone can give rise to lateralized activity resembling a CDA, it does not reproduce the full pattern of sign reversals observed in Experiment 1. The absence of a polarity flip in Condition 3, together with the weaker overall evidence for Model 1, suggests that the robust CDA reversals in Experiment 1 cannot be explained solely by attentional selection.

## Discussion

To gain insight into the neurophysiological processes underlying real-time manipulations of working memory representations in a dynamic world, the current study examined the CDA – an electrophysiological marker of WM load – in a novel experimental paradigm where subjects were required to encode objects (colored discs) into WM and then update the spatial location associated with them. In a separate experiment, subjects completed a visually identical task without having to remember the colors. Together, the results suggested that dynamic updates to the relative location of objects held in mind are reflected in qualitative changes in the CDA. These changes were related to both the updated WM information and attentional processes.

In Experiment 1, we observed that the CDA followed the mental movement in real time, reversing its sign when the objects moved into the opposite hemifield, and then again when they returned to the original hemifield. These results were consistent with our main hypothesis that mentally updating the location of a memorized item causes a cortical transfer of the neural representation of that item, although it should be noted that the evidence for Condition 5 was relatively weak. Interestingly, our findings stand in contrast to those of an earlier study that found no CDA sign reversal when objects were encoded in one hemifield and later tested in the opposite one^26^. We suspect the effect observed here requires a task with strong emphasis on updating contents in real time to prevent the updating during the test period itself. Consistently, including subjects with poor accuracy on the behavioral task reduced the effect (Supplemental Figure 2). However, the source of the effect could also be the continuous visuospatial attention required in our task, if CDA is driven by attention rather than WM. The aim of Experiment 2 was to examine to what extent attentional tracking can explain the effect.

The results from Experiment 2 suggested that neither interpretation alone could fully account for the dynamically updated CDA. If the CDA effects observed in Experiment 1 were solely due to attentional tracking of moving placeholders, then similar CDA patterns should have emerged even in the absence of a WM requirement. Conversely, if the CDA strictly reflected maintained and manipulated WM content, no sustained lateralized activity would be expected when subjects attended to the placeholders without encoding or storing color information^17,27^. Instead, the observed CDA response in Experiment 2 showed partial activation, suggesting that attentional tracking alone may elicit some CDA-like activity, but the full effect seen in Experiment 1 likely depends on the presence of active WM representations. Crucially, unlike in Experiment 1, the observed CDA did not reverse polarity in any condition at any time point, indicating that polarity reversals could be a distinctive marker of active updating of spatial WM information.

Another interesting observation was the strong negative post-movement CDA in Condition 5 of the attention task. This strong negative CDA, compared with the relatively weak and inconclusive effect in the same condition in the WM task, represents a notable divergence between the two experiments. One tentative explanation is that this difference relates to the absence of a polarity reversal in Condition 3 of the attention task. If attentional tracking remained anchored to the original hemifield even when both discs crossed the midline, then when they returned in Condition 5 the CDA would re-emerge strongly negative, unlike in the WM task where intermediate remapping attenuated the effect. Further work will be needed to clarify the mechanisms behind the divergence in Condition 5.

Previous research has shown CDA to reflect filtering (tracking the number of relevant items, and not distractors^14,21^) and to dynamically track various transformations of the WM load – such as grouping^28,29^, splitting^30^, and sequential encoding^17^, – but its response to spatial manipulations, up until very recently^32^, has been explored only in the context of multiple-object tracking (MOT)^24^. Notably, Drew et al.^24^ observed CDA reversing polarity in their MOT task, which they refer to as ‘handoff of attention’. However, the lack of such a reversal in our second experiment contradicts this purely ‘attentional’ interpretation. Additionally, previous findings suggest that object-tracking related CDA fades approximately 350 milliseconds after the objects stop moving^23^, whereas we observed sustained CDA throughout the post-movement delay periods with objects held fixed, suggesting WM-rather than attention-related activity. What is more, a recent study by Wang et al.^31^ suggests that the classic MOT tasks may include an implicit WM demand by requiring participants to maintain the spatial representations of the tracked stimuli in WM, meaning that past observations from MOT might have reflected more than just attentional processes too.

In both WM and MOT literature, CDA has widely been considered to reflect item-based processes, be it WM maintenance or attentional tracking. Recently, a refined view has emerged, suggesting it to reflect a common process of assigning spatiotemporal ‘pointers‘^33^. In a combined WM and MOT task, Styrkowiec et al.^32^ observed CDA reflecting tracking load rather than WM load when both were simultaneously tested. The authors interpreted this as evidence that CDA reflects an indexing mechanism rather than content-related activity. However, this view cannot account for the observed differences between our two experiments. Even if the two disks in Experiment 2 were assigned a single pointer (which could explain a weaker pre-movement CDA compared to Experiment 1), it would not explain the lack of polarity reversal during or after the movement. Moreover, the pointer-based accounts fail to explain the growing body of evidence showing CDA to reflect more than discrete item-based processing^34^, as it has been observed to be sensitive to perceptual complexity^35,36^, semantics^37,38^, and familiarity^39^. Our results seem in line with these less discrete accounts of the nature of CDA.

In conclusion, our results suggest that the cortical location of activity related to both attention and WM was updated to meet behavioral demands as the spatial location of remembered objects changed. This would put them in line with single neuron activity recorded from non-human primates suggesting that neural information retaining WM object identity is transferred across hemispheres as item locations are updated^12^. However, since the nature of the CDA is still debated, whether the observed activity reflects pointers (or a similar indexing mechanism) or the WM content itself remains an open question. This could be further tested by using methods directly decoding object identity from EEG^40^.

## Methods

### Subjects

A total of 26 (Experiment 1) and 31 (Experiment 2) subjects were recruited via Karolinska Institutet’s online recruitment system. All subjects provided informed written consent before the start of the experiment and were reimbursed 100 SEK per hour. The study was approved by the Swedish Ethical Review Authority and was performed according to the Declaration of Helsinki. All subjects reported normal or corrected-to-normal visual acuity and no neurological disorders. Prior to participation in Experiment 1, color vision was tested using the short (6-plate) version of the Ishihara’s Test for Color Deficiency^41^. The test was considered passed if all plates were read normally. All subjects passed the test.

### Experiment 1

Data from one subject was excluded from analysis due to excessive noise in the EEG and data from an additional six subjects were excluded due to poor behavioral performance (see below for details). The analysis reported in the main text was performed on the data from the remaining 19 subjects (M_age_ = 28.2, SD_age_ = 5.2; 11 female, 8 male; 3 left-handed), who each performed 480 trials across 5 conditions (see below). According to a recent meta-analysis^42^, this was sufficient to test our main hypothesis. We verified that our conclusions did not critically depend on the behavior-based participant exclusions (Supplemental Material).

### Experiment 2

One subject was excluded from analysis due to not completing the full experimental session. Two subjects were excluded due to excessive noise in the EEG. Additionally, two more were excluded due to poor behavioral performance. The final sample size was 26 subjects (M_age_ = 28.2, SD_age_ = 5.1; 14 female, 12 male; 2 left-handed, 1 ambidextrous). As in Experiment 1, all subjects performed 480 trials across 5 conditions.

### Stimuli and procedure

All tasks were completed on a Windows PC with a Dell E2422H monitor (24”, 1920 × 1080 pixels, 60 Hz refresh rate). The viewing distance was approximately 58 cm, maintained by positioning subjects’ chin on a chin rest. The tasks and stimuli were generated and displayed using MATLAB with the Psychophysics Toolbox Version 3 (PTB-3) extension^43–45^. During the experiment, the subjects’ pupil size and eye movements were recorded using a Gazepoint GP3 eye tracker. Trials in which participants broke fixation by moving their gaze were automatically aborted, but the experimental session continued until they had successfully completed a total of 480 trials. A break in fixation was defined as the average gaze position straying more than 5.4° from the center of the screen.

Experiments 1 and 2 were structurally and visually identical, except for the format of the probe. The experimental session consisted of two parts, with a 5-10-minute rest period in between. Each part consisted of 240 trials, divided into 12 blocks of 20 trials each. Blocks were separated by 10-second breaks. Before the start of the experimental session, subjects completed a training block of 20 trials. EEG was not recorded during the training and the behavioral data were not included in the analysis.

On each trial subjects were presented with an array of four colored discs on a gray (RGB = 128 128 128) background. The colors of the discs were chosen from an isoluminant color wheel (CIE L*a*b* color space, [L = 70, a = −6, b = 14, radius = 49]), discretized to 180 colors and calibrated to the testing monitor. The colors were drawn from a uniform distribution on the color wheel but with the constraint that no two colors were separated by less than 30 degrees. In Experiment 1, subjects were asked to memorize two of the four presented colors, located in a particular hemifield. The relevant hemifield was randomized across trials and indicated by a pre-cue. In Experiment 2, subjects were instructed to ignore the colors.

### Trial structure

A trial started with a fixation period, during which a fixation point (two concentric circles: black, radius of 0.3°, and white, radius of 0.1°) was presented at the center of the screen. The subjects’ focus on the fixation point was monitored by the eye tracker to ensure 300 ms of uninterrupted fixation, meaning that moving their gaze away from the screen center extended the fixation period rather than aborting the trial. Breaking fixation at any later point, from the appearance of the cue and until the appearance of the probe, however, aborted the trial. A cue in the form of an arrow, indicating the to-be-attended side, was presented for 250 ms following the fixation period. Then, after a variable delay of 300-400 ms with only the fixation point visible, four colored discs (radius=1.3°) were presented for 500 ms, located at the corners of an invisible square centered on the fixation point. The distance between the fixation point and each stimulus was 8.1°. This was followed by a delay period of 1000 ms, during which only four identical black outlines of the discs remained on the screen, with their colors having disappeared at the start of the delay. Then, these outlines started moving along an invisible orbit centered on the fixation point, in identical direction and with identical speed, so that both the distances between the discs and the distance between the discs and the fixation point remained the same. The direction and angle of the movement was randomly selected from the range of [0°, 360°] in steps of 30°, meaning there were 13 possible angles in total. The movement was performed at a constant speed of 180°/s, meaning that its duration depended on the angle, with the shortest movement lasting 0 s (angle of 0°) and the longest movement lasting 2 s (angle of 360°). Another delay followed the movement, during which the discs’ outlines remained visible in their new locations. The duration of the delay was set to 3 s minus the duration of the movement, so that the total duration of movement and delay was always exactly 3 s, of which at least 1 s was delay. After the delay, in Experiment 1, the outlines of the discs disappeared and a memory probe – a color wheel – appeared at the new location of one of the previously cued discs. Subjects were asked to recall the color of that disc by picking it on the color wheel. Subjects had unlimited time to respond. After their response was submitted, they received feedback showing both the color they had chosen and the correct color for 500 ms, which concluded the trial. In Experiment 2, all outlines stayed on the screen, with one – the probe – changing its color to white. Subjects were asked to indicate whether the probe was one of the two discs they were tracking by pressing the “F” or “L” key. The keys were labeled with the words “YES” and “NO”, with the label assignment randomized between participants. The probe was a tracked disc in exactly half of the trials. After the response was submitted, the fixation point changed color to reflect whether it was correct (green) or incorrect (red). This feedback was shown for 500 ms. The intertrial interval was 1000 ms.

### Conditions

We defined five experimental conditions, based on the final locations of the two cued discs: Condition 1 (0°, 30°), in which both discs remained in the original hemifield; Condition 2 (60°, 90°, 120°), in which one of the discs transferred to the other hemifield; Condition 3 (150°, 180°, 210°), in which both discs transferred to the other hemifield; Condition 4 (240°, 270°, 300°), in which both discs transferred to the other hemifield and then one of them returned to the original hemifield; and Condition 5 (330°, 360°), in which both discs transferred to the other hemifield and then returned to the original one.

### Trial balancing

The attended hemifields were set to have equal numbers of trials (240) and within each attended hemifield, the numbers of trials per movement condition were likewise set to be equal (48). The numbers of trials per angle within each condition were balanced as well (24 or 16 for conditions with 2 or 3 angles, respectively). The sign of the angle, which determined whether the movement would be clockwise or counterclockwise, and the probed item were randomized across trials without enforcing equal numbers. Trial order was randomized, separately for each subject.

### Participant exclusion based on poor behavioral accuracy

A pivotal aspect of our task design for Experiment 1 was that accurate performance was possible only if subjects mentally updated locations of stimuli held in WM as the disc outlines moved on the screen (see main text). Subjects with large response errors likely failed to perform this mental updating. Since our main interest was to study neural correlates of mental updating, data of such subjects were expected to be uninformative, essentially only adding noise. Therefore, we excluded subjects with unusually large errors from the main analyses. To this end, we imposed a performance criterion that excluded all subjects with an average response error of at least two standard deviations above the group mean, which was enforced iteratively until no subjects fulfilled this criterion. This resulted in a total of 6 exclusions (compare Figure 2a with Supplemental Figure 3 to see the performance of the excluded subjects relative to the rest). To make sure that our main findings do not critically depend on this exclusion, we reran the analysis with these subjects included. As anticipated, the results were highly similar, albeit with reduced statistical strength due to a lowered signal-to-noise ratio in the EEG data (see Supplemental Figure 2), suggesting that accurate performance was indeed pivotal. For Experiment 2, the same criterion was applied but only once, rather than iteratively, leading to 2 exclusions.

### Behavioral analysis

#### Experiment 1

For each participant we calculated the mean response error for each of the 13 motion angles. We calculated this error both with respect to the color of the target item and the color of the non-target (the cued but not probed) item. To assess how these two types of errors related to the motion angle, we used a Pearson correlation test, applied to participant-averaged errors (hence, 19 data points per correlation test). To estimate how the swap rate and memory precision related to the motion angle, we fitted an equal-precision model with non-target responses^20^ to the raw response data, separately for each participant. In this model, the response error on each trial was drawn from a Von Mises distribution with a concentration parameter *κ*. Moreover, there was a certain probability, *p*_swap_, that the response was based on the memory of the non-target item instead of the target. To investigate potential effects of motion angle *α* on memory precision and swap rate, we modelled both these variables as linear functions, through *κ*(*α*) = *a* + *bα* and *p*_swap_(*α*) = *c* + *dα*, where *a, b, c* and *d* were fitted as free parameters. The best-fitting parameters for each participant were found through maximum-likelihood estimation, that is, by finding the values of *a, b, c*, and *d* that maximized

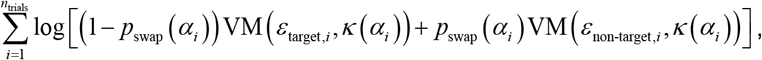

where VM(*ε, κ*) is a zero-mean Von Mises distribution with concentration parameter *κ*, evaluated at *ε*; *ε*_target,*i*_ and *ε*_non-target,*i*_ denote the response errors relative to the target and non-target item, respectively, on trial *i*; *α*_*i*_ denotes the motion angle on trial *i*. The optimization was performed in Python, using the Bayesian Adaptive Direct Search (BADS) package^46^.

#### Experiment 2

Since the purpose of the task in Experiment 2 was to ensure that participants were actively tracking rather than passively viewing the movement, we did not perform any behavioral analyses apart from computing the performance accuracy defined as the ratio of correct responses to the total number of trials.

#### EEG recording

EEG was acquired at 2048 Hz from 64 active Ag/AgCl electrodes using BioSemi ActiveTwo system. The electrodes were placed on an elastic cap according to the International 10-20 system. Two additional electrodes were placed on the left and right mastoids. The vertical electrooculogram (VEOG), used to detect eye blinks, was recorded from electrodes located 2 cm above and below the right eye. The horizontal electrooculogram (HEOG), used to measure horizontal eye movements, was recorded from electrodes located 1 cm lateral to the external canthi.

### EEG preprocessing

Preprocessing and analysis of the EEG data was carried out using MNE-Python^47^. For the initial visual inspection, all electrodes were re-referenced offline to the average of the mastoids. If faulty electrodes were (manually) detected during or after the recording, they were interpolated using the MNE interpolate_bads() function with default parameters. This was done for 2 out of 190 (10 electrodes x 19 participants) electrodes kept in the main CDA analysis for Experiment 1, and for 9 out of 260 electrodes for Experiment 2. As excessive noise was discovered in the mastoid electrodes of several participants, EEG data were then re-referenced to the average of all 64 electrodes. The continuous data were bandpass-filtered using a finite impulse response (FIR) filter to 0.1-30 Hz and down-sampled to 256 Hz. A Picard^48^ Independent Component Analysis (ICA), as implemented in MNE, was then run on the filtered and down-sampled data and the obtained components were automatically tested (using MNE function find_bads_eog() that utilizes correlation with EOG channels) and manually inspected in order to detect and remove ocular artifacts (blinks and horizontal eye movements). The results of automatic detection aided the process, but the final decision was made based on manual inspection. EEG was epoched between -200 and 4500 ms time-locked to stimulus onset. Each epoch, therefore, included 200 ms of pre-stimulus fixation, 500 ms of stimulus presentation, 1000 ms of pre-movement delay, 0-2000 ms of stimuli movement, and 1000-3000 ms of post-movement delay.

Automatic trial rejection was used to remove trials contaminated with muscle artifacts, channels blocking, slow drifts, and step-like voltage changes in the electrodes used for the ERP analyses. The detection algorithms and rejection criteria were implemented as described in^49^. The remaining trials were manually inspected to make sure no obvious artifacts had been missed. To aid reproducibility, manual rejection was substituted with an additional criterion of peak-to-peak amplitudes exceeding 150 μV, as the result of applying it was virtually identical. Subjects with fewer than 360 (75%) clean trials were excluded from analyses. Thus, for the remaining 19 subjects in Experiment 1, we rejected on average 1.6% (SD = 1.4%) of trials, resulting in an average of 472 trials per participant. For the 26 subjects in Experiment 2, an average of 2.4% (SD = 3.1%) trials were rejected, leaving an average of 468 clean trials per participant.

### EEG analysis

#### CDA extraction

Artifact-free epochs were baseline-corrected using the 200 ms of pre-stimulus fixation as baseline and trial-averaged separately for left and right attended sides. All ERP analysis was restricted to the 10 posterior channels reported to show maximal CDA in memory tasks^50^: P5/P6, P7/P8, PO3/PO4, PO7/PO8, and O1/O2. The difference (CDA) waves were computed per participant, by subtracting their average activity recorded at ipsilateral sites from the average activity recorded at contralateral sites, with laterality defined relative to the attended (cued) side.

For statistical analysis, we measured CDA as the mean amplitude between 300 and 1000 ms post stimulus offset for pre-movement delay (analyzed in the Supplemental Material) and the mean amplitude between 300 and 1000 ms post movement end per condition in post-movement delay (main analysis). The movement end per condition was defined as the time point at which the longest movement included in the condition ended, e.g. 30° movement for Condition 1 and 210° movement for Condition 3.

#### Frequentist t-tests

To test the predictions of our main model against the data, we performed Student’s t-tests on the time-averaged CDA measures. The null hypothesis was specified as H_0_: *μ*=0. The model predicted H_0_ to be true in Conditions 2 and 4 and false in Conditions 1, 3, and 5. Since the model predicted a specific effect direction in each of the latter three conditions, we used one-tailed tests for those conditions. For the two conditions where the null hypothesis was predicted to be true, we used a two-tailed test.

#### Bayesian t-tests

A limitation of frequentist tests is that they can only provide evidence *against* the null hypothesis. A high *p*-value means that the data do not provide evidence against the null hypothesis, but it would be a statistical fallacy to interpret such a finding as evidence *in favor* of the null^51^. Therefore, we also analyzed the data using Bayesian t-tests. This type of test quantifies evidence by means of a Bayes Factor (BF), expressing how strongly the data support one hypothesis relative to another hypothesis. BFs above 1 provide support for a given hypothesis, below 1 evidence against. We use the interpretation scale from ^52^ to map Bayes factors to verbal labels such as ‘weak’ and ‘strong’ evidence. An advantage of BFs is that they can quantify evidence both for and against any well-defined hypotheses, including the null^53^. In the Bayesian evaluation of the model predictions, we computed for each condition the BF for the hypothesis following from our main model, denoted H_M_, relative to an alternative hypothesis H_A_, as listed in Table 1. Note that in Condition 1 we compared the model prediction, H_M_: *μ*<0, only against “*μ*=0”, not against “*μ*>0”. Our motivation for this was that published literature on CDA consistently reports the amplitude to be negative or non-existent and rarely or never as positive. Therefore, “*μ*>0” has near-zero prior plausibility and should, therefore, not be included when doing Bayesian testing (including it would have increased the evidence for our model, so the reported value can be considered as a conservative estimate). For the other conditions we had no strong prior expectations, because there is no prior literature on the effect of mental updating on CDA. Therefore, we specified the alternative hypothesis in those conditions as the complement of the model’s hypothesis (e.g., when the model predicts “>0”, the alternative would be defined as “≤0”). We computed BFs using the Bayesian t-test in JASP^54^, which allows to test four different types of hypotheses (*μ*<0, *μ*=0, *μ*>0, *μ*≠0) against each other. A limitation is that it does not allow to directly test the “compound” alternative hypotheses that we specified for Conditions 3 and 5. Instead, we computed these BFs as the average BF obtained from testing H_M_ separately against each component in the compound hypothesis. Hence, in Condition 3, we took the average of the BF obtained from testing H_M_: μ>0 against H_A_: μ=0 and the BF obtained from testing H_M_: μ>0 against H_A_: μ<0. The latter BF (for H_M_: μ>0 against H_A_: μ<0) could also not be computed directly in JASP. Instead, we computed it as the ratio of BF_+0_ and BF_−0_: 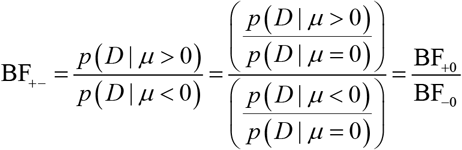. The BF for Condition 5 was computed in a similar way. In all Bayesian t-tests, the default prior was used (a Cauchy distribution with a scale parameter of .707).

**Table 1.** Overview of hypotheses tested in the Bayesian t-tests. For each condition, we calculated a Bayes Factor BF_MA_, quantifying the evidence in favor of the hypothesis H_M_ relative to H_A_. Bayes Factors larger than 1 indicate support for the model prediction, while Bayes Factors smaller than 1 indicate evidence against the model prediction.

| Condition | $H_M$ | $H_A$ | Computation of $BF_{MA}$ |
| --- | --- | --- | --- |
| 1 | $\mu < 0$ | $\mu = 0$ | $BF_{MA} = BF_{-0}$ |
| 2 | $\mu = 0$ | $\mu \neq 0$ | $BF_{MA} = BF_{01}$ |
| 3 | $\mu > 0$ | $\mu \leq 0$ | $BF_{MA} = \frac{1}{2}BF_{+0} + \frac{1}{2}\frac{BF_{+0}}{BF_{-0}}$ |
| 4 | $\mu = 0$ | $\mu \neq 0$ | $BF_{MA} = BF_{01}$ |
| 5 | $\mu < 0$ | $\mu \geq 0$ | $BF_{MA} = \frac{1}{2}BF_{-0} + \frac{1}{2}\frac{BF_{-0}}{BF_{+0}}$ |

### Bayesian mixed-effects modeling

To evaluate the three competing hypotheses about CDA dynamics on all 5 conditions at once, we fitted Bayesian mixed-effects models to post-movement CDA amplitudes (300 – 1000 ms). Analyses were performed in Python using the *bambi* package with MCMC sampling (4 chains, 2000 draws after 1000 warm-up, target acceptance 0.9). CDA values were expressed in microvolts, and all models included subject-level random intercepts; random slopes were added when condition-specific effects were estimated.

We tested three models that aligned with the theoretical accounts. In all three models, the intercept was fixed to 0, which is motivated by the baselining procedure. In Model 1 (hemispheric transfer), the beta coefficients for Conditions 1 and 5 were fitted as free parameters with a negative prior; the beta coefficient for Condition 3 was fitted as a free parameter with a positive prior; and the beta coefficients in Conditions 2 and 4 were fixed to 0. In Model 2 (uniform CDA), a single beta coefficient with negative prior was fitted across the mean across all five conditions. In Model 3, a beta coefficient with negative prior was fitted to the first condition, while the coefficients for the other four conditions were fixed to 0.

All priors were Half-Normal distributions with σ = 1.5. Model fits were compared using leave-one-out cross-validation (LOO) with pseudo-BMA model weights; WAIC was used when LOO was unstable.

*Robustness checks:* To verify that our statistical results did not critically depend on the specific time windows used in the CDA calculations, we also ran the analyses with time windows locked to the movement end for each individual motion angle (resulting in 13 separate time windows instead of 5) and with a single window that was time-locked to the end of post-movement delay. The results (see Supplemental Figure 1 and text) were consistent with the results reported in the main text.

## Data availability

All preprocessed data will be made available upon publication.

## Code availability

All code will be made available upon publication.

## Acknowledgements

This work was funded by ERC starting grant 949131 and Swedish research council project grant VR 2022-02328.

