## Supplemental Material for "Contralateral delay activity during dynamic spatial updates in working memory"

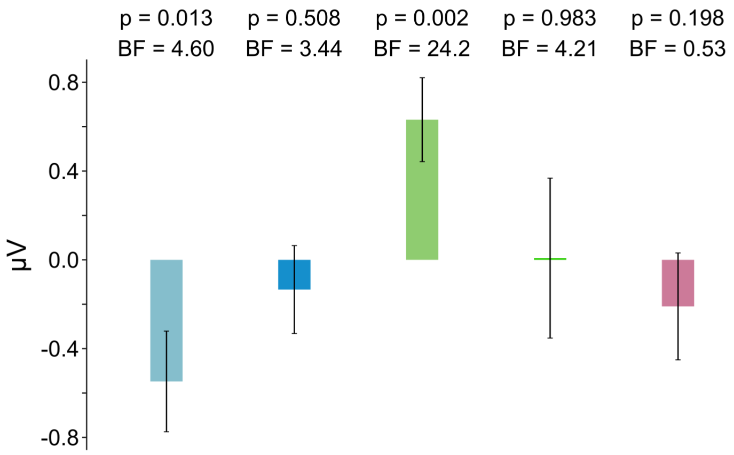


**Supplemental Figure 1.** Same as Figure 3c, but with average CDA amplitude obtained from time windows locked to 300-1000 ms post movement end rather than time locked to condition end. Error bars denote SEM. The p-values were obtained from t-tests against the null hypothesis of CDA being 0. The BFs were obtained from Bayesian t-tests and quantify the evidence in favor of the predictions made by Model 1 (see Methods).

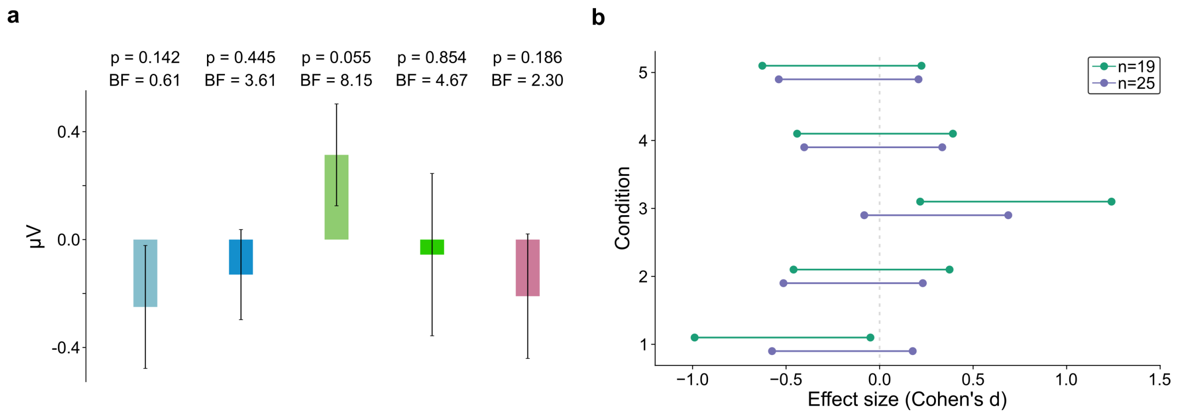


**Supplemental Figure 2. a.** Same as Figure 3c, but for 25 subjects (including the ones excluded from main analysis based on behavior). **b.** Bayesian 95% credible intervals for CDA amplitude, split by condition and presented separately for the cases with behavior-based exclusions (n=19, as in the main text; green) and without such exclusions (n=25; purple). The CDAs were taken from the time periods indicated in Figure 3b.

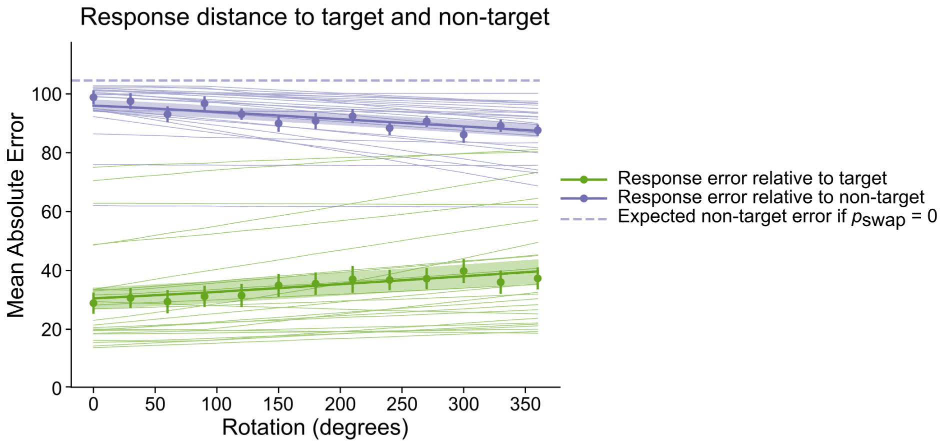


**Supplemental Figure 3.** Same as Figure 2a, but for 25 subjects (including the ones excluded from main analysis based on behavior).

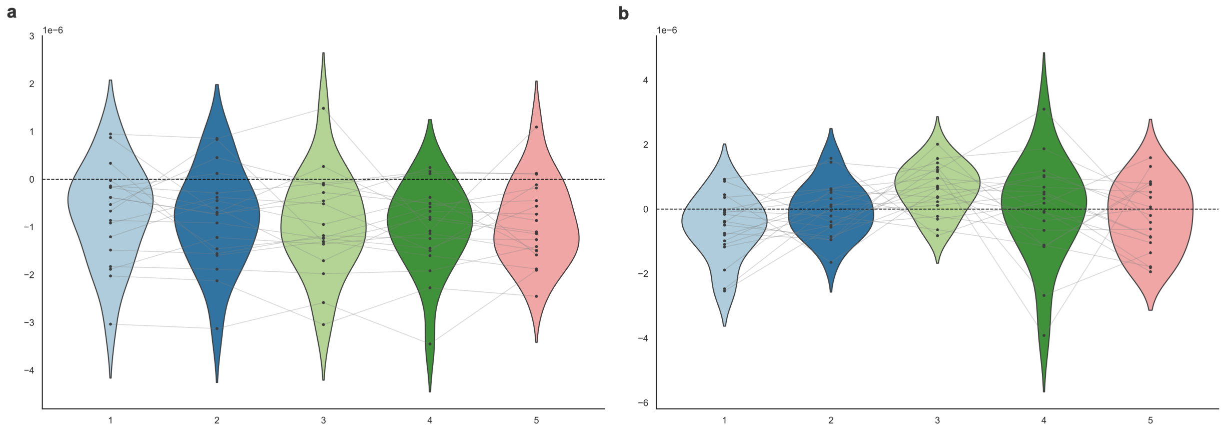


**Supplemental Figure 4.** Distribution of average CDA amplitudes per condition in Experiment 1. **a.** Before the movement start (300-1000 ms post stimulus offset). **b.** After the movement (same time window as in Figure 3c). Black dots represent average CDA of individual participants, with light gray lines connecting data from the same participant across conditions.

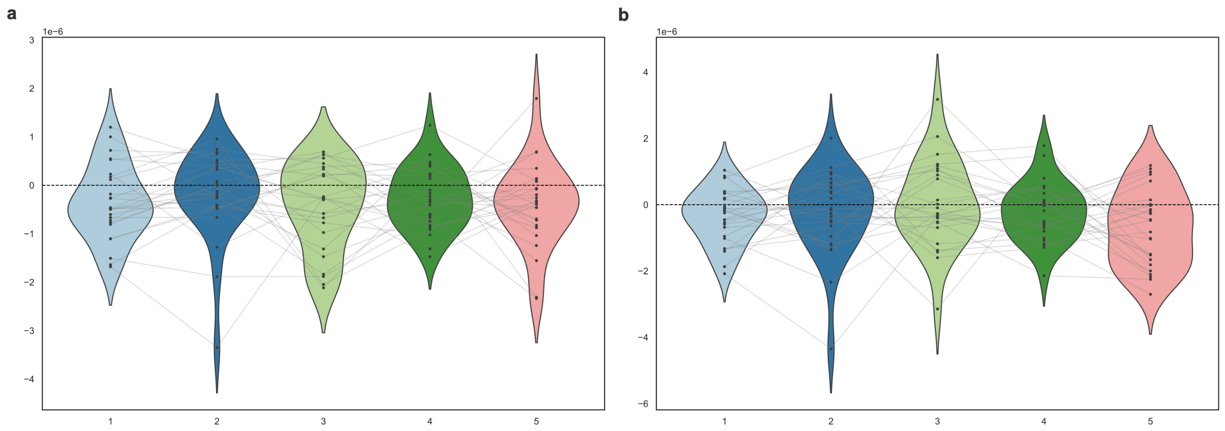


**Supplemental Figure 5.** Distribution of average CDA amplitudes per condition in Experiment 2. **a.** Before the movement start (300-1000 ms post stimulus offset). **b.** After the movement (same time window as in Figure 3e). Black dots represent average CDA of individual participants, with light gray lines connecting data from the same participant across conditions.

### *Supplemental text:*

We additionally performed Bayesian t-tests to test Model 1 in the same way as presented in Figures 3c and 3e, but instead with CDA windows time-locked to the end of post-movement delay (last 700 ms) rather than to the movement end. For Experiment 1, the BFs for this time window per condition were: condition 1: BF = *3.30* ("<0" vs "=0"); condition 2: BF = *3.97* ("=0" vs "≠0"); condition 3: BF = 4.67 (">0" vs "≤0"); condition 4: BF = *3.83* ("=0" vs "≠0"); condition 5: BF = 0.60 ("<0" vs "≥0"). For Experiment 2: condition 1: BF = 1.76; condition 2: BF = 4.06; condition 3: BF = 0.29; condition 4: BF = 2.56; condition 5: BF = 20.2.
